# Miniprotein inhibitors of the *Staphylococcus aureus* efflux transporter NorA

**DOI:** 10.64898/2026.03.05.709893

**Authors:** Priyanka Mishra, Adam Chazin-Gray, Gaëlle Lamon, David Kim, David Baker, Nathaniel J. Traaseth

## Abstract

Multidrug efflux pumps transport antibiotics across the cellular membrane resulting in resistance conferred to the host organism. Efflux pump inhibitors (EPIs) potentiate the efficacy of antibiotics by blocking drug efflux and hold promise as adjuvant therapeutics in the fight against multidrug resistant pathogenic bacteria. A hurdle in the field has been the lack of selectivity of small molecule EPIs which often display off-target toxicity due to non-specific binding. To tackle this specificity challenge, we aimed to maximize an inhibitor’s binding surface area to efflux pumps by designing miniprotein EPIs using computational protein design and an *E. coli* co-expression assay to screen inhibition in cells. We used *S. aureus* NorA as a model efflux transporter since it confers drug resistance to fluoroquinolones, puromycin, and other cytotoxic compounds. Starting from a focused miniprotein library of only 86 members, we identified inhibitors in the screen that blocked NorA transport under active efflux conditions *in vitro*. Our most promising inhibitor I-23 was validated by solving a cryo-EM structure of the miniprotein in complex with NorA, which stabilized the transporter in the outward-open conformation. I-23 has a ferredoxin-like fold with one of its β-hairpins inserted into the substrate binding pocket of NorA and other parts of the globular fold occupying the shallow pocket and making extensive intermolecular contacts with NorA. An arginine residue on the tip of the hairpin loop was positioned near an anionic patch required for NorA antibiotic efflux. The identified structural motifs in this work could be employed to explore the molecular properties of peptidoglycan penetration; full realization of the therapeutic potential of the designed miniprotein inhibitors will require determining the principles for facilitating passage of ∼7 to 8 kDa miniproteins across the peptidoglycan bacterial cell wall.

## Introduction

Multidrug resistant *Staphylococcus aureus* strains, such as methicillin-resistant *S. aureus* or MRSA, are Gram-positive bacterial pathogens that contribute to treatment failures, increased morbidity and mortality among patients, and an economic burden to healthcare systems ^1^. Many strains display resistance to the major antibiotic classes and evade the immune response, leading to severe infections such as sepsis, pneumonia, meningitis, and endocarditis ^2, 3^. Antibiotic efflux is a key resistance mechanism that decreases the intracellular concentration of antibiotics, allowing bacteria to survive in the presence of these toxic compounds ^4–7^. Inhibitors of efflux have potential as therapeutics by sensitizing the organism to antibiotics which are otherwise ineffective due to this resistance mechanism ^8^.

Antibiotic transport from the cytoplasm to periplasm requires efflux pumps to transition between at least two conformations, inward- and outward-facing states ^9–11^. The main inhibition strategies involve direct competition with the antibiotic binding site using small molecules or trapping a single conformation in the catalytic cycle. Despite promising preclinical research, there are no approved efflux pump inhibitors (EPIs) on the market for use in humans. Some clinical trials of EPIs were halted due to the acute toxicity of small molecules, which likely stemmed from their off-target effects ^12–14^. The source of non-specific binding may arise from the hydrophobic character of the efflux pump substrate binding pocket and the likelihood that small molecules match this hydrophobicity.

Increasing the surface area of the inhibitor-transporter interface may improve affinity and reduce off-target binding. A larger binding interface could also be an effective strategy for trapping the conformation of the protein and bypassing the need for competition with substrates. A few observations suggest transporters can accommodate larger molecules within the substrate binding pocket, including the presence of native and nonnative lipids used for structure determination and the presence synthetic antibody loops used as crystallization chaperones in X-ray crystallography or fiducial marks in cryo-EM ^8, 15, 16^. In some cases, the deepest parts of the substrate binding pocket are occupied by these molecules and further accompanied by interactions in the shallower parts of the substrate binding cleft near the lipid-water interface. Given these observations, we hypothesized that miniproteins would serve as efficient scaffolds to occupy both the deep and shallow pockets with the goal of maximizing the inhibitor-transporter surface area. Miniproteins are small, folded proteins spanning ∼1-10 kDa that have emerged as promising next-generation therapeutics because they combine the pharmacokinetics of peptides and stability of proteins ^17–19^. Miniprotein-based therapeutics have achieved antibody-like specificity and affinity while mitigating the limitations of larger scaffolds ^20–27^.

In this work, we developed miniprotein inhibitors of *S. aureus* NorA, a secondary active multidrug efflux transporter from the major facilitator superfamily (MFS) that confers resistance to antibiotics, biocides, and cationic dyes ^28–30^. NorA overexpression is a common attribute of MRSA ^31^ and blocking its function with a selective EPI may rejuvenate the activity of currently existing ineffective antibiotics ^32, 33^. The first structures of NorA solved in complex with antigen-binding fragments (Fabs) trapped NorA in the outward-open conformation where the CDRH3 loop was inserted into the substrate binding pocket ^8^. Despite the inhibitory binding interaction, Fabs are too large (∼50 kDa) to effectively penetrate the peptidoglycan layer of *S. aureus* ^34^. Therefore, to reduce the inhibitor scaffold while retaining a large binding surface area, we designed miniproteins using computational approaches and screened their inhibitory potential using a bacterial co-expression experiment under active efflux conditions. Initiating a screen from a small library led to the identification of miniprotein inhibitors that were validated using binding assays and structural experiments.

## Results

### Design and identification of miniprotein inhibitors to NorA

Miniprotein designs were initiated by generating scaffolds for the CDRH3 loop of Fab36 bound to NorA (PDB ID: 7LO8) using RFdiffusion ^35^ or a novel PyRosetta-based loop grafting approach called Autograft (**Figure 1**). Sequences for the backbone of each scaffold were assigned while either fixing the Fab36 CDRH3 loop sequence or separately allowing ProteinMPNN ^36^ to redesign the entire miniprotein sequence including that of the loop motif. The complexes of each set of designs were predicted with AlphaFold2 (AF2) Initial Guess ^37, 38^ while templating NorA in the outward-open conformation and filtered by their predicted alignment errors and confidence scores. The final miniprotein designs were obtained by resampling these filtered designs using partial diffusion, followed by ProteinMPNN and AF2 Initial Guess, and finally filtering predictions using stricter alignment error and confidence score cutoffs.

**Figure 1.**
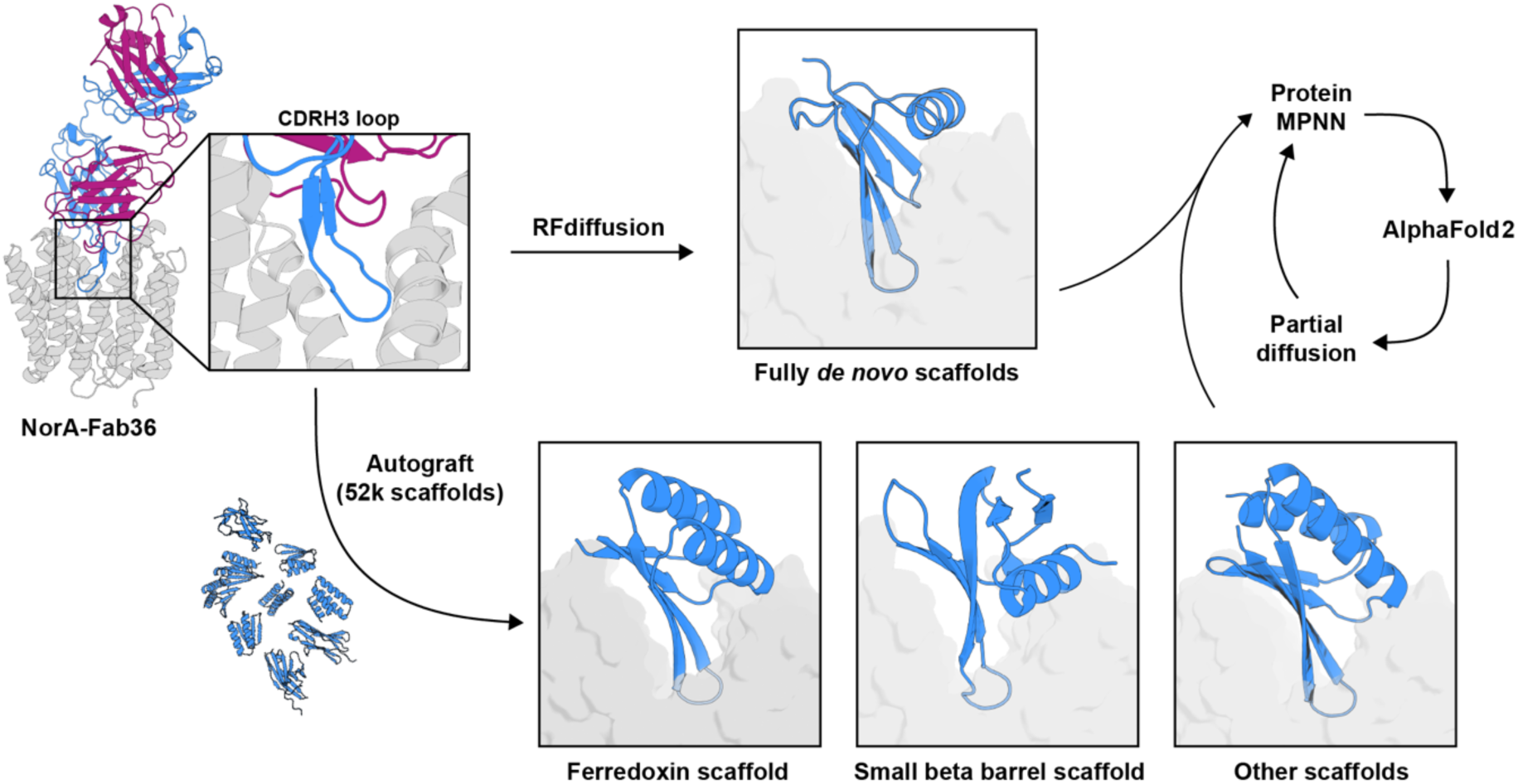
Computational design strategy of miniprotein binders to NorA. Design of miniproteins scaffolding the CDRH3loop of Fab36 bound to NorA (PDB ID: 7LO8) using RFdiffusion or Autograft. The sequences of each miniprotein backbone were assigned by ProteinMPNN while either fixing the Fab36 CDRH3 loop sequence or separately allowing ProteinMPNN to redesign the entire miniprotein sequence including CDRH3. The structure of the complex was then predicted with AlphaFold2 (AF2) Initial Guess. Final designs were obtained after resampling using partial diffusion, followed by ProteinMPNN sequence design, and AF2 Initial Guess predictions using stricter alignment error and confidence score cutoffs.

A total of 86 miniprotein sequences were selected for screening using an *E. coli* co-expression assay with the native *acrB* gene disrupted by a transposon insertion. This assay screens for inhibition under efflux active conditions following transformation of bacteria with two separate plasmids (**Figure 2a**). The first plasmid encodes *norA* on a pTrc His2C vector with a *trc* promoter that drives gene expression in a leaky manner while the second encodes miniproteins on a pBAD33 vector with an arabinose inducible araBAD promoter. Each miniprotein gene contained a signal peptide to direct the protein to the periplasm since miniproteins were designed to target the outward-facing conformation of NorA which is accessible from the periplasm. Inhibition of NorA blocks efflux and sensitizes bacteria to norfloxacin resulting in growth inhibition, while a lack of inhibition enables NorA to confer resistance to this antibiotic resulting in no growth inhibition (**Figure 2a**). To ensure reproducible results that could be compared quantitatively across samples, three control growth inhibition experiments were also performed: (*i*) NorA expressed with miniproteins in the absence of norfloxacin to identify when a miniprotein displayed toxicity to the organism independent of NorA inhibition, (*ii*) NorA expressed without miniproteins in the presence of norfloxacin to assess maximal resistance, and (*iii*) inactive NorA (E222A dead mutant ^8, 39^) expressed without miniproteins and in the presence of norfloxacin to assess background resistance of the *E. coli* strain.

**Figure 2.**
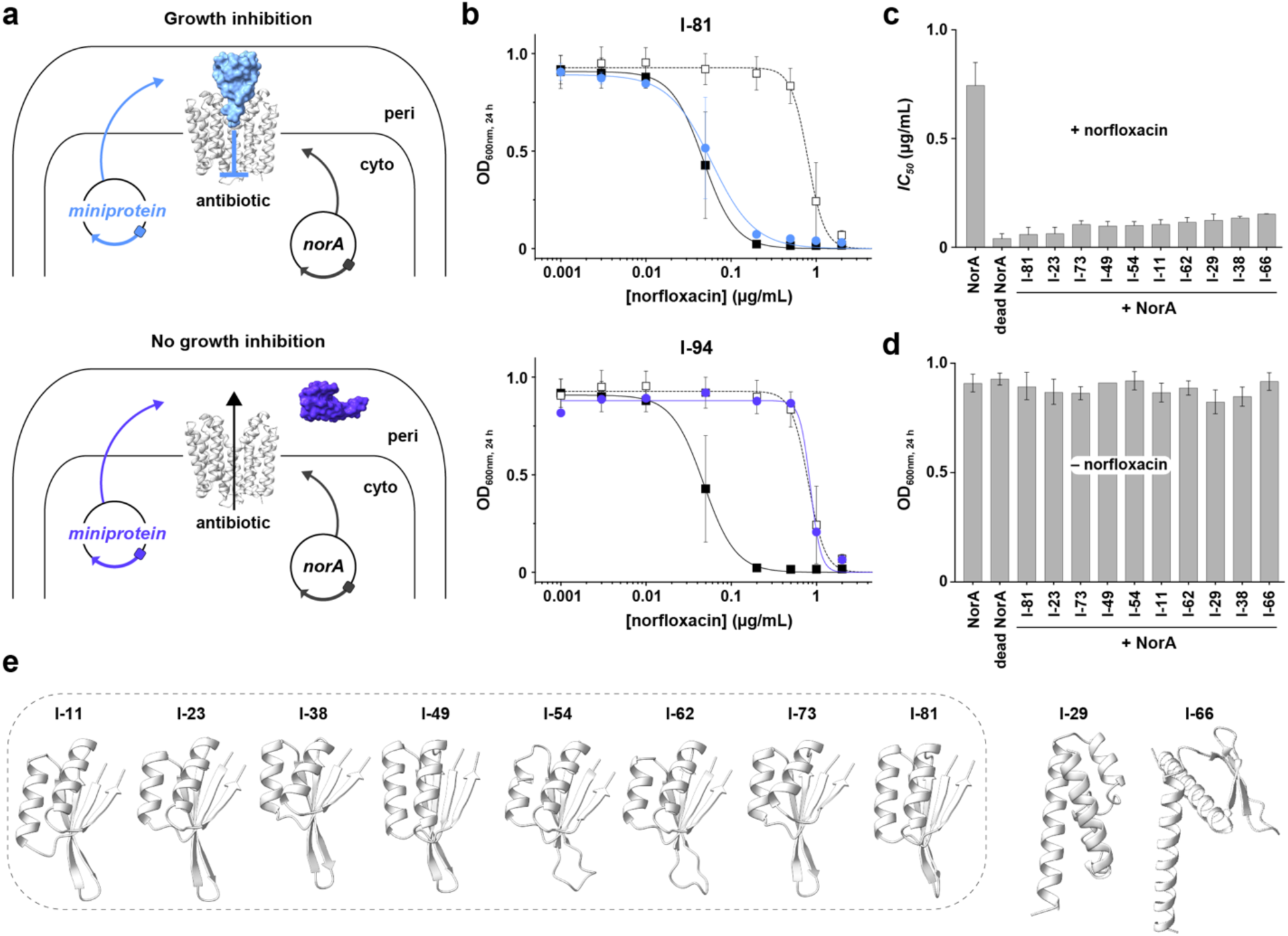
*E. coli* co-expression experiments identify miniprotein inhibitors of NorA. **a.** Schematic of *E. coli* co-expression experiment where NorA and miniprotein genes are encoded on different vectors. NorA was cloned in the pTrcHis2C vector that yields constitutive expression and each miniprotein was cloned in the pBAD33 vector that is arabinose inducible. The miniproteins were secreted to the periplasm with a signal peptide on the N-terminus. The top and bottom schematics display the presence or absence of NorA inhibition by miniproteins, respectively. **b.** Example growth inhibition results for determining the *IC_50_* values to norfloxacin in *E. coli* DH5α 11*acrB*. These co-expression experiments were performed in the presence of 1.6% arabinose. In each panel, the solid black squares correspond to NorA, the open black squares are dead NorA (E222A mutant), and the blue or purple circles are miniproteins I-81 (top) or I-94 (bottom). The fitted curves are used to derive the *IC_50_* values. These data reveal I-81 as an inhibitory miniprotein and I-94 as a non-inhibitory miniprotein. **c.** *IC_50_* values of norfloxacin growth inhibition experiments in *E. coli* DH5α 11*acrB* corresponding to the top miniprotein inhibitor hits expressed in the presence of NorA. *IC_50_* values of norfloxacin growth inhibition experiments for NorA and dead NorA (E222A mutant) in the absence of miniproteins are included for reference. **d.** OD_600nm_ values at 24 h for *E. coli* DH5α 11*acrB* co-expressing each of the top miniprotein inhibitors and NorA in the absence of norfloxacin. The OD_600nm_ values for strains expressing NorA and dead NorA (E222A mutant) but no miniproteins are included for reference. **e.** AF2 predicted structures of the top 10 miniprotein hits in the absence of NorA. The dotted box indicates miniproteins possessing a ferredoxin-like fold.

Growth was quantified by measuring the optical density at 600 nm over 24 hours for NorA co-expressed with each miniprotein. Each set of growth inhibition experiments were fitted to obtain the half maximal inhibitory concentration (*IC_50_*) and further converted into a percent NorA inhibition based on NorA and the inactive NorA mutant controls (**Figure S1a; Table S1**). We discovered seven miniproteins that displayed > 90% NorA inhibition, 15 miniproteins with 80 to 90% inhibition, and 12 miniproteins with 50 to 80% inhibition. The best inhibitors displayed *IC_50_* values and growth inhibition curves nearly indistinguishable from those of the dead mutant control indicating maximal inhibition (e.g., I-81 and I-23), while miniproteins that failed to inhibit or express resembled experiments with the NorA control (e.g., I-94) (**Figure 1b, c**). Notably, *E. coli* expressing the top miniprotein hits in the absence of norfloxacin grew to the same optical density as the control samples, which indicated an antibiotic adjuvant role for the miniproteins where growth inhibition was not due to toxicity or off-target effects (**Figure 1d**).

To validate miniprotein binding, pull down experiments were performed by co-expressing NorA with the top 10 miniproteins that each possessed a C-terminal FLAG tag. Growth inhibition experiments revealed that the FLAG tag did not alter NorA inhibition by miniproteins (**Figure S1b**). NorA was purified by isolating the membrane fraction, solubilizing the transporter with LMNG detergent, and passing the lysate over a metal affinity chromatography column to bind the C-terminal His-tag on NorA. The eluted fractions from the column were immunoblotted for the presence of the miniprotein’s FLAG tag ^40–43^. Immunoblotting experiments showed that purified NorA was bound to each of the 10 miniproteins (**Figure S1c**). These results indicated that inhibition observed in co-expression assays stemmed from direct binding and suggested sufficiently strong binding to co-purify throughout the NorA purification.

### Cryo-EM structure validation of miniprotein binding

AF2 structural predictions of the top miniprotein hits revealed that eight out of 10 adopted a ferredoxin-like fold when bound or unbound to NorA (I-11, I-23, I-38, I-49, I-54, I-62, I-73, and I-81) (**Figure 1e**). Of these eight, five shared the CDRH3 sequence from Fab36, which was used as an input motif for the miniprotein design process (**Table S2**). Given the similar predicted ferredoxin-like fold of several of the hits and the sequence identity with Fab36, we selected I-23 for validation of binding and structural predictions since it was the inhibitor with the lowest *IC_50_* that shared no sequence identity with the previously developed Fab. I-23 was expressed, purified, and used in a size-exclusion chromatography (SEC) binding assay. We observed that I-23 co-migrated with NorA in SEC experiments, indicating sufficiently good binding to form a stable complex throughout the assay (**Figure S2a**).

To determine the cryo-EM structure of this complex, we used a NorA-BRIL fusion protein ^44^ bound by the BRIL-recognizing BAG2 antibody ^45^ to increase the size of the complex and to serve as a fiducial mark during data processing. SEC-purified complexes containing I-23 were frozen on cryo-EM grids, screened for suitable ice thickness, and used to acquire a large cryo-EM dataset. Data processing resulted in a cryo-EM map with an overall map resolution of 3.08 Å and showed the expected proteins in the complex, including I-23, NorA, BRIL, and BAG2 (**Figure S2b**). To further improve map quality for I-23, a focused refinement was performed, which yielded the final cryo-EM map with a resolution of 2.79 Å (**Figure 3a**). The local map resolution ranged from 2.5 to 3.3 Å with most of the interfacial regions between NorA and I-23 displaying resolutions of 2.5 to 2.9 Å (**Figure S3**). Given the quality of the maps, a structural model of the NorA-I-23 complex was constructed (**Figure 3b; Figure S4; Table S3**).

**Figure 3.**
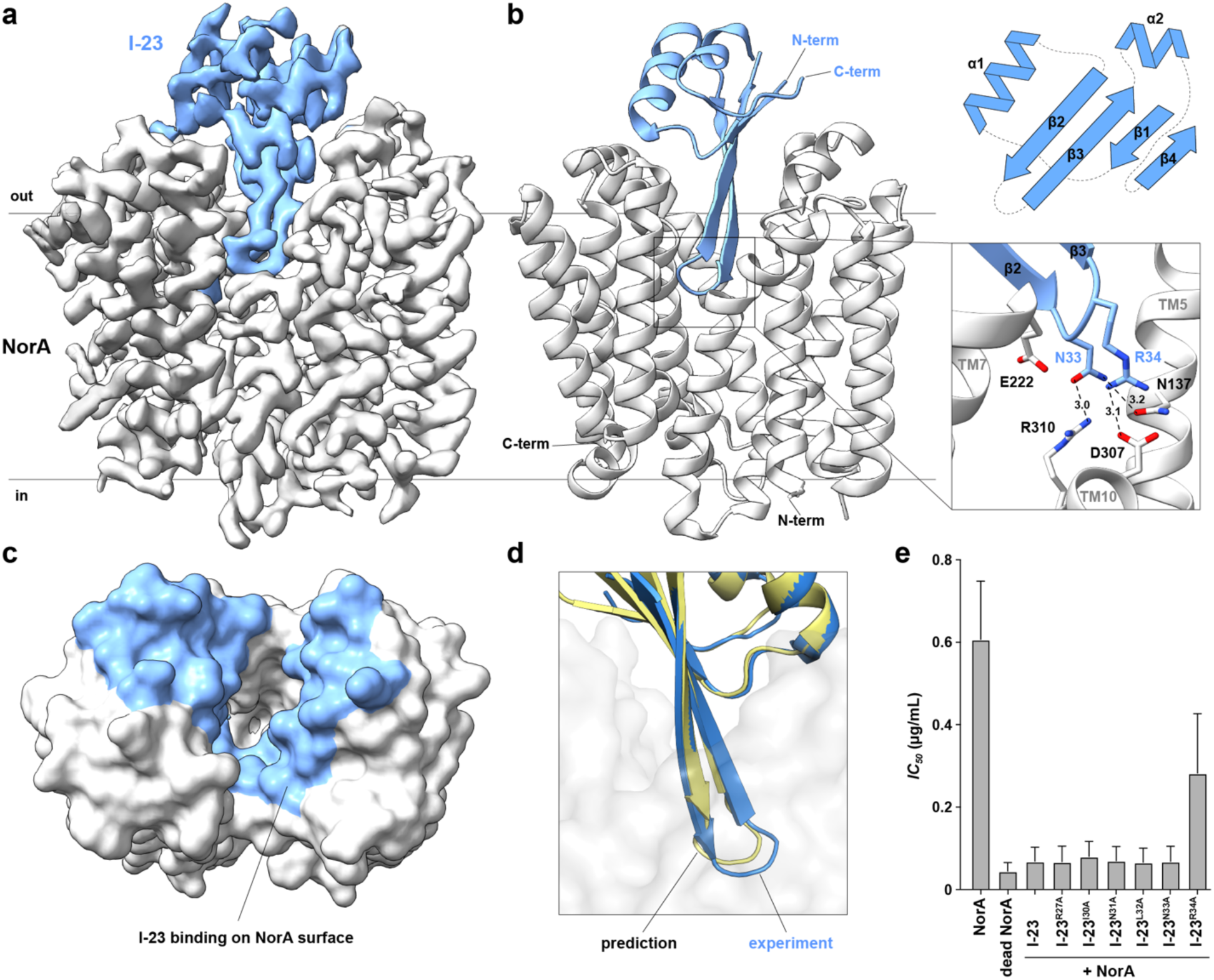
Cryo-EM structure of NorA bound to the miniprotein inhibitor I-23. **a.** Cryo-EM map of the NorA-I-23 complex obtained after local refinement. NorA is displayed in grey and I-23 is shown in light blue. **b.** Structural model of the NorA-I-23 complex (left) with an expanded view of the β-hairpin loop (bottom right) and a topology view of I-23 (top right). NorA and I-23 are displayed in grey and light blue cartoon representations, respectively. Dotted lines indicate distances between heavy atoms (displayed in Å). **c.** Surface view representation of NorA looking into the substrate binding pocket from the periplasmic direction. Light blue colors indicate the region I-23 is within 5 Å from NorA residues. **d.** Superimposition of the I-23 AF2 predicted model (yellow) onto the experimental I-23 structure (light blue). The experimental NorA structural model is displayed in a transparent grey surface representation. **e.** *IC*_50_ values for norfloxacin derived from E. coli growth inhibition experiments of NorA expressed in the presence of I-23 and the indicated mutants. *IC*_50_ values were derived by fitting the *OD*_600nm_ values from the 24 h time points. *IC*_50_ values for NorA and dead NorA (E222A mutant) in the absence of miniproteins are included for reference. Data are presented as mean values ± s.d. among three independent experiments.

The structure showed NorA in the outward-open conformation with I-23 adopting a ferredoxin-like fold with topology βaββaβ (**Figure 3b**). Interacting residues from I-23 covered ∼1,400 Å^2^ of surface area within NorA (**Figure 3c**). The central two β-strands of I-23 formed a hairpin loop within the substrate binding pocket of NorA, where one β-strand spanned from residues Lys24 to Leu32 and the other from Ile35 to Thr43. The I-23 loop residues of Asn33 and Arg34 were most buried in the central cavity with Asn33 in proximity to NorA residues Asn137 and Arg310 for making hydrogen bonds and Arg34 near NorA residue Asp307 for making a salt bridge (**Figure 3b**). There were also several intramolecular interactions within NorA bridging these residues, including nearby interactions among Arg310 with Glu222 and Asp307 which likely played a role in stabilizing interactions with Asn33 from the loop of I-23. The position of the β-hairpin of I-23 closely matched the position of the CDRH3 loop of Fab36, albeit with only a single shared Arg residue on the tip of the loop (**Figure S5a**).

A comparison of the NorA-I-23 experimental structure with the AF2 Initial Guess predicted structure showed a backbone RMSD of 0.681 Å (**Figure S5b**). Note that this structural comparison displayed only a portion of NorA which was constrained to the outward-open conformation to expedite computational design and predictions. Despite the similarity between the experimental and predicted structures, there was a notable deviation for the tip of the β-hairpin loop backbone which was displaced by ∼2 Å from the predicted structure and for the side chains of Asn33 and Arg34 which differed by 2 to 4 Å from the prediction (**Figure 3d**). Nevertheless, the cryo-EM structure serves as validation of our approach to discover miniprotein inhibitors using computational protein design and the functional screen.

### A key arginine residue in I-23 is present in the miniprotein’s β-hairpin

Based on our structural model, we mutated six interfacial residues of the β-hairpin within I-23 to assess their importance for NorA inhibition. Alanine mutations were introduced at R27, I30, N31, L32, N33, and R34 since these positions were in contact with NorA residues within the substrate binding pocket in our model (**Figure 3b**). Growth inhibition assays in *E. coli* were performed on these I-23 mutants as in the original screen, followed by quantification of norfloxacin *IC_50_* values for each mutant co-expressed with wild-type NorA. Of these mutants, only the R34A mutant displayed a loss of inhibition relative to the parent I-23 miniprotein (**Figure 3e**). This result indicated that the key residue in the miniprotein is the cationic arginine within the β-hairpin loop that binds to the acidic region of the NorA substrate binding pocket. We cannot rule out that other mutants may reduce the binding affinity to NorA because the concentration of miniproteins in the assay may exceed the binding constant. Nevertheless, based on these results, we conclude that the key factor for binding is the positively charged arginine which is positioned to make a salt bridge with Asp307 of NorA.

## Discussion

In this work, we designed a small miniprotein inhibitor library to NorA, identified inhibitors using a functional screen, and validated our first-generation miniprotein inhibitor (I-23) by determining a cryo-EM structure of I-23 bound to NorA. The ferredoxin-like fold of I-23 provides a scaffold for stabilizing the β-hairpin that inserts into the pocket of NorA. The hairpin loop possesses an asparagine and arginine residue pair which form electrostatic interactions with the acidic patch on NorA composed of Glu222 and Asp307. These interactions lock NorA in the outward-facing conformation by preventing conformational exchange, which is required for proton-coupled antibiotic transport ^39^. Comparison of NorA bound to I-23 and NorA bound to Fab25 or Fab36 indicates that the key intermolecular interactions stemming from two different folds engage the substrate binding pocket through the same anionic patch at Glu222 and Asp307, a critical location for inhibition ^8^. In contrast to the Fabs, the I-23 miniprotein (7.8 kDa) uses ∼34% of its entire surface area to make intermolecular contacts with NorA, which is significantly higher than Fab25 and Fab36 which use ∼6% and ∼10% of their surface area, respectively. This improved inhibitor efficiency and ability to contour portions of the ferredoxin-like scaffold into the upper cleft of the substrate binding pocket offers a way to further optimize contacts which can likely be extended to other transporters.

Despite identifying I-23 as an *in vitro* binder and inhibitor to NorA in our *E. coli* assay, preliminary growth inhibition assays with I-23 added to MRSA cultures in the presence of norfloxacin failed to inhibit bacterial growth (**Figure S6**). Possible reasons are that I-23 failed to efficiently penetrate through the peptidoglycan (PG) layer of *S. aureus* due to its size or surface charge. Prior work showed that the PG layer is porous with an estimated minimum diameter of 2-3 nm (∼25 kDa) and progressively smaller pores near the lipid bilayer ^34^. We hypothesize that miniproteins too hydrophobic will not possess sufficient solubility to penetrate the hydrophilic PG layer and those too charged may enter but interact too strongly with the PG layer resulting in attenuated diffusion. Given the small size of I-23 (7.8 kDa), it is possible that the surface charge facilitates non-specific binding and influences penetration through the PG layer. Therefore, redesigning the surface charge of this miniprotein might yield improved penetration and result in MRSA growth inhibition in the presence of norfloxacin. Further optimization of these miniproteins, such as those with a ferredoxin-like fold and others, will enable this hypothesis to be tested, leading to additional insight into the physical properties governing protein penetration through the PG layer of *S. aureus*.

In conclusion, our study demonstrates the first designed miniprotein targeting a membrane protein multidrug transporter. This work establishes a generalizable design framework that can be applied to scaffold other antibody loops onto small protein folds. I-23 is likely to serve as a useful protein scaffold for developing miniprotein inhibitors to other MFS-type multidrug transporters from clinically relevant bacterial pathogens. A question emerging from this work is how to design small proteins that can efficiently penetrate the PG layer. Our design strategy and identification of the ferredoxin-like fold will serve as a tool for answering this question with implications for the design of biotherapeutic antibiotic adjuvants.

## Materials and Methods

### Computational design of miniproteins

Miniprotein backbones were generated in two ways. First, 10,000 backbone designs were generated by scaffolding the Fab36 CDRH3 loop from the NorA-Fab36 cryo-EM structure (residues Y128 to W139 from chain H of PDB ID: 7LO8 ^8^) using RFdiffusion ^35^. Additionally, other Fab36 loop scaffolds were generated using a PyRosetta-based loop grafting approach called Autograft that staples an input motif onto an existing miniprotein scaffold library (see “Autograft” section below). Two approaches were taken for sequence design of these miniprotein backbones. 10 sequences for each backbone was assigned by ProteinMPNN by *(i)* fixing the Fab36 CDRH3 loop sequence (YYYAWRVGG) and *(ii)* separately allowing ProteinMPNN to redesign the entire miniprotein sequence including the loop itself. The complexes of both sets of designs were then predicted with AF2 Initial Guess ^38^ while templating NorA in its outward-open conformation. Designs with predicted aligned error (pAE) scores under 15 and predicted local distance difference test (pLDDT) scores over 70 were then resampled using partial diffusion, followed by ProteinMPNN sequence redesign, and AF2 structure prediction as before. Final designs were selected using AF2 Initial Guess metrics for the complex (pLDDT_binder ≥ 80, pAE_interaction ≤ 10), and AF2 monomer metrics for the minibinder alone (pLDDT_monomer ≥ 80, Ca_RMSD < 1).

### Autograft

*De novo* miniprotein scaffolds were grafted onto the hairpin loop of the heavy chain CDRH3 from the NorA-Fab36 complex structure (PDB ID: 7LO8) using Autograft (https://github.com/davidekim/autograft), a python script that utilizes PyRosetta to identify adjacent β-strand pair segments of three residues in the CDRH3 hairpin loop that have minimal RMSD to β-strand pair segments in each scaffold. The segment pair from the hairpin loop was selected based on the number of C_β_ atom contacts (≤ 8 Å) that the central residue of each segment had with the target (≤ 5) and the total number of contacts (≥ 10). If the RMSD was ≤ 1.5 Å with matching β-strand pleating between the superimposed segments, the scaffold was grafted at the midpoint residue position of each segment using the superimposed coordinates. If there were no clashing backbone atoms in the grafted complex, the structure was idealized using PyRosetta. Only idealized grafts were used for partial diffusion refinement and sequence design.

### Construction of E. coli DH5α 11acrB mutant

The *acrB* gene was removed from the DH5α *E. coli* genome using a recombineering protocol and a temperature sensitive *pSIJ8* plasmid ^46, 47^. A single colony transformed with pSIJ8 was grown overnight in 5 mL 2YT media supplemented with carbenicillin at 30 °C. A 100-fold dilution of overnight culture was prepared in 50 mL 2YT media without carbenicillin and induced with 10 mM arabinose for growth at 30 °C until the OD_600nm_ value reached 0.4-0.6. An uninduced culture was used as a negative control. The culture was washed several times in sterile distilled water, and the final pellet was resuspended in 10% glycerol for the electroporation of the PCR product containing a kanamycin cassette and the 50 bp gene homology arms ^48^. The recovered cells were plated on LB agar plates supplemented with carbenicillin and kanamycin for growth at 30 °C overnight. Single colonies were grown in 20 µL for 2 h and the presence of the kanamycin cassette was verified with PCR using 1 µL of culture. To remove the antibiotic cassettes, the remaining culture was grown for 6 h in the presence of 50 μM rhamnose followed by plating on LB agar plates supplemented with carbenicillin at 30 °C. Single colonies were verified by PCR for the excision of the kanamycin cassette. PCR verified colonies were grown at 42 °C to facilitate curing of the temperature sensitive pSIJ8 plasmid.

### Cloning of miniprotein library in pBAD33

The miniprotein genes were ordered from IDT as double stranded eBlocks in a 96-well plate format. Primers were designed to amplify the genes as PCR products suitable for cloning into pBAD33 (forward primer: TATGCAGCCCTCGAGT; reverse primer: GCAACGTGAAGCTTCC). The genes were PCR amplified and digested with restriction enzymes (XhoI and HindIII) for ligation in the double digested pBAD33 plasmid with gentamicin resistance ^49^ using standard cloning procedure.

### Inhibition assays of NorA by miniproteins in E. coli

NorA was cloned in a pTrc His2C vector (gift of David Hooper ^28^) while the miniprotein library was cloned in a pBAD33 vector. Co-transformation experiments were performed using chemically competent DH5α Δ*acrB E. coli* by simultaneously adding vector encoding wild-type NorA and the miniprotein. For the co-transformation of controls, empty pBAD33 with no miniprotein gene was used with wild-type NorA (WT, positive control) or E222A NorA (dead mutant, negative control). The 86 miniproteins in pBAD33 and NorA in the pTrc His2C vector were co-transformed in separate experiments to test the inhibitory potential of individual miniprotein with NorA. The selection of co-transformed cells were performed by plating the cells on LB agar plates infused with gentamicin and carbenicillin after 1 h of recovery from the heat shock. A single bacterial colony from the transformation was inoculated into 5 mL TBG media in the presence of gentamicin and carbenicillin. The final concentration of gentamicin and carbenicillin used in LB agar plates and TBG medium was 50 and 100 μg/mL, respectively. The culture was grown for ∼18 h at 30 °C at a shaking speed of 200 rpm. Overnight cultures were diluted by 300 to 500-fold into fresh TBG medium containing gentamicin, carbenicillin, norfloxacin (10, 5, 2, 1, 0.5, 0.2, 0.05, 0.01, 0.003 and 0.001 μg/mL), and 1.6% arabinose. The OD_600nm_ for each culture was normalized before diluting the cells to have a similar OD_600nm_ after the dilution. Growth curves were acquired by measuring the OD_600nm_ every 15 min using a BioScreen Pro C device at 30 °C over a period of 24 h with moderate shaking. Growth data was collected for all 86 miniproteins in the presence of wild-type NorA and by using the wild-type NorA and dead mutant as controls in each experimental setup.

### Pull down experiments of miniproteins

DNA corresponding to the FLAG-tag sequence (DYKDDDDK) was added to the pBAD33 vector using PCR (forward primer: ATCATACTCGAGGCATAGGCATTTG, reverse primer: GGTCATAAAGCTTAGGAGGAGGTGGATCAGATTACAAGGACGATGATGACAAGTAAGTCGACCTGCAGGCATG). Miniprotein genes for the top 10 hits were cloned into the modified pBAD33 (pBAD33_FLAG) and tested for inhibition of NorA in *E. coli*, as described in the section above. For pull down experiments, several bacterial colonies with the vector expressing NorA and the individual miniprotein hits (I-23, I-62, I-49, I-54, I-11, I-29, I-81, I-73, I-38 and I-66) were inoculated into 50 mL of TBG medium in the presence of 0.4% arabinose, gentamicin and carbenicillin. The inoculated medium was grown at 30 °C, 200 rpm for 20 h. The cells from each culture were collected by centrifugation at 5,000 rpm using a Beckman centrifuge equipped with a JLA 8.1 rotor (Beckman Coulter) for 20 min at 4 °C. The cell pellets were stored at −80 °C.

Cell pellets were thawed, resuspended and lysed via homogenization in 40 mM Tris (pH 8.0), 400 mM NaCl, and 10% glycerol. The lysate was clarified by centrifugation at 13,000 x g for 30 min at 4 °C using a Beckman centrifuge equipped with a JA-25.50 rotor (Beckman Coulter). The supernatant was centrifuged for 4 h at 35,000 rpm to collect the membrane fraction using a Beckman centrifuge equipped with a Type 45 Ti fixed-angle rotor (Beckman Coulter). Membranes were resuspended and solubilized in 20 mM Tris pH 8.0, 300 mM NaCl, 10 mM imidazole, 10% glycerol, and 2% (w/v) lauryl maltose neopentyl glycol (LMNG; Anatrace) for 2 h while stirring at 4 °C in glass vials. After 2 h, the solubilized membrane fraction was centrifuged at 15,000 rpm for 20 min at 4 °C to remove the insoluble membrane fraction. The supernatant was mixed with 200 μL of pre-equilibrated HisPur Ni-NTA Resin (Thermo Fisher Scientific) and incubated with gentle stirring at 4 °C for 30 min. NorA was bound to Ni-NTA affinity resin and washed with 20 mM Tris pH 8.0, 200 mM NaCl, 10% glycerol, 0.2% (w/v) LMNG) and 25 mM imidazole (wash buffer) to remove unbound proteins followed by elution with wash buffer containing 400 mM imidazole. Immunoblotting analyses were performed on purified NorA using primary antibodies diluted at 1:2,000 for MYC (Genscript) and FLAG-tag (Genscript) present at the C-termini of NorA and miniprotein, respectively. Immunoblotting analyses were repeated two independent times.

### I-23 expression and purification

I-23 was cloned in a pMAL C2X expression vector (New England Biolabs) and transformed into BL21(DE3) T1R *E. coli*. Maltose binding protein (MBP) was fused to the N-terminus of I-23 and expressed in the cytoplasm (i.e., the C2X vector contains no secretion peptide). For protein expression, transformed cells were inoculated in LB media and grown at 37 °C at a shaking speed of 200 rpm. At an OD_600nm_ of ∼1.0, protein expression was induced by addition of 1 mM IPTG (final concentration). Following IPTG addition, cells were grown at 20 °C for 16-18 h at a shaking speed of 200 rpm. Cells were harvested by centrifugation at 5,000 rpm using a Beckman centrifuge equipped with a JLA 8.1 rotor (Beckman Coulter) and stored at −80 °C until further use. Cell pellets were thawed, resuspended in 20 mM Na_2_HPO_4_ (pH 7.3) and 300 mM NaCl (purification buffer) and supplemented with 1 mg/mL hen egg lysozyme (Sigma), EDTA, glycerol, protease inhibitors. Following homogenization, the lysate was centrifuged at 20,000 rpm for 20 min at 4 °C in a Beckman centrifuge equipped with a JA-25.50 rotor (Beckman Coulter). The clarified supernatant was loaded onto a column with pre-equilibrated amylose resins (E8021L, NEB). The column was washed with purification buffer to remove non-binding proteins and the miniprotein was eluted in the purification buffer containing maltose. The MBP-miniprotein fusion protein was incubated overnight with tobacco etch virus (TEV) protease at 4 °C followed by passing the cleaved proteins over a Nickel affinity column to remove uncleaved fusion protein, MBP, and TEV protease. The cleaved miniprotein was recovered in the flow through followed by the final purification using size exclusion chromatography (SEC) with a Superdex 75 Increase 10/300 GL (Cytiva) preequilibrated with purification buffer.

### Fab expression and purification

The BAG2 gene ^45^, rigidified in the elbow region of the heavy chain ^50^, was purchased (Twist Bioscience), subcloned into a pTac expression vector, and transformed into the 55244 strain of *E. coli* (ATCC), as previously described ^44^. BAG2 was expressed and purified following the procedure we previously reported ^44^ with the following modifications. Briefly, the Fab was expressed through leaky expression by incubating of *E. coli* cells for 24 h at 30 °C in TBG media with moderate shaking. Cells were harvested by centrifugation at 6,000 x g (JLA 8.1 rotor, Beckman Coulter) and stored at −80 °C until further use. Cell pellets were thawed, resuspended in phosphate buffer (20 mM Na_2_HPO_4_ pH 7.0) supplemented with 1 mg/mL hen egg lysozyme (Sigma) and homogenized. After centrifuging at 27,000 x g for 1.5 h at 4 °C in a Beckman centrifuge equipped with a JA-25.50 rotor (Beckman Coulter). The clarified supernatant was then filtered and loaded onto a protein G column (Cytiva). The column was extensively washed with phosphate buffer to remove unbound protein and the targeted Fab was eluted with 100 mM glycine (pH 2.7) and immediately neutralized with 2 M Tris buffer (pH 8).

### NorA-BRIL expression and purification

The pET29b vector containing the NorA-BRIL gene with a three alanine linker was transformed into C43 (DE3) *E. coli* and autoinduced with essentially the same procedure previously reported ^44^. Cells were harvested by centrifugation and stored at −80 °C until further use. Cell pellets were thawed, resuspended, and homogenized in lysis buffer (40 mM Tris pH 8.0, 400 mM NaCl, 10% glycerol), and centrifuged at 13,000 x g for 30 min at 4 °C in a Beckman centrifuge equipped with a JA-25.50 rotor (Beckman Coulter). The membrane fraction was isolated by centrifuging for 2.5 h at 186,000 x g using a Beckman ultracentrifuge equipped with a Type 45 Ti fixed-angle rotor (Beckman Coulter) and solubilized in 20 mM Tris pH 8.0, 300 mM NaCl, 10% glycerol (v/v), 10 mM imidazole, and 1% (w/v) LMNG (Anatrace) for 2 h. The solubilized protein was incubated with nickel affinity resin (Thermo Fisher Scientific) at 4 °C for 1 h to maximize the binding to the target protein. Unbound proteins were washed away with low-imidazole buffer (20 mM Tris pH 8.0, 200 mM NaCl, 10% glycerol (v/v), 0.2% (w/v) LMNG, 25 mM imidazole). NorA-BRIL protein was eluted with high imidazole buffer (20 mM Tris pH 7.5, 200 mM NaCl, 10% glycerol (v/v), 0.2% (w/v) LMNG, 400 mM imidazole). Subsequently, the carboxy-terminal decahistidine tag in the elution fractions was removed from NorA-BRIL by incubation with TEV protease at while dialyzing for 16 h at 4 °C (20 mM Tris pH 7.5 and 100 mM NaCl). The TEV protease was captured by affinity chromatography against nickel resin while the cleaved NorA-BRIL protein recovered in the flowthrough. PMAL-C8 amphipol (Anatrace) was added to the flow through fractions at a 5/1 ratio (w/w) of PMAL-C8/NorA-BRIL and incubated with stirring for 16 h at 4 °C. LMNG was removed from the sample by addition of Bio-beads (Bio-Rad) (10 g for each 50 mL of protein) and incubated with the sample for 24 h at 4 °C. The monomeric protein was finally purified by SEC using a Superdex 200 10/300 column (Cytiva) preequilibrated with buffer set to pH 5.0 (20 mM Na_2_HPO_4_ pH 5.0 and 100 mM NaCl). Peak fractions of the target protein were pooled and stored at 4 °C until further use.

### Cryo-EM sample preparation and screening

NorA-BRIL/BAG2/I-23 (1:3:3 mole:mole ratio) sample was prepared by adjusting the pH of each protein to 7.5 separately, mixing them together and adjusting the NaCl concentration to 300 mM. The complex was purified with SEC using a Superdex 200 10/300 column (Cytiva) preequilibrated with buffer (20 mM Tris pH 7.5 and 100 mM NaCl). The most concentrated fraction (50 μM) was collected and used for freezing cryo-EM grids.

To prepare cryo-EM grids, 4 μL of sample was applied onto UltrAuFoil 300-mesh R1.2/1.3 grids (Quantifoil) glow-discharged with the PELCO easiGlow GlowDischarge system (Ted Pella Inc.). Grids were blotted on both sides for 3.5 or 4 s at 100% humidity and a temperature of 16 °C and then plunge-frozen into liquid ethane using a FEI Vitrobot Mark IV. The grids were screened on a Talos Arctica microscope (200 kV X-FEG electron source, two condenser lens system) equipped with a Gatan K3 camera (5760 × 4092).

### Cryo-EM data collection

Cryo-EM data for NorA-BRIL/BAG2/I-23 sample was acquired on a Titan EF-Krios microscope at 105,000x nominal magnification under 300 kV (imaging system: Gatan Bioquantum K3 camera) ^51–53^. Zero-loss images were taken using a GIF-Quantum energy filter with a 20-eV slit. Leginon software version 3.6 was used for automated data collection ^54^. Movies were collected with a physical pixel size of 0.4125 Å, sample received an accumulated dose of 49.586 e^-^/Å^2^ and the ice thickness on-the-fly was measured to select the holes with the desired ice thickness for data collection ^52^. A total of 9,026 movies were collected at a nominal defocus range of −0.7 – 2.4 μm ^55, 56^. On-the-fly movie processing was executed using MotionCor2 software version 1.5 and CTFFIND4 software version 4.1.13, under regulatory purview of Appion ^57^. Meanwhile, on-the-fly particle picking was performed using Warp ^58^.

### Cryo-EM image processing and map analysis

The NorA-BRIL-BAG2-I-23 dataset was processed in cryoSPARC (version 4.4.0) ^59^. The contrast transfer function was estimated for the dose-weighted micrographs. Initial 2D class averages and *ab-initio* model reconstitution were generated using well-resolved 2D classes generated from particles picked by Warp ^58^. Topaz training ^60^ was used for additional rounds of particle picking and generated a total of 6.7 million selected particles. This set of particles was then refined using heterogenous 3D refinement. Particles from classes corresponding to complete NorA-BRIL-BAG2-I-23 complexes were retained for further rounds of *ab initio* model building and heterogeneous 3D classification. After multiple rounds, a non-uniform 3D refinement step was used to generate the final 3.08 Å resolution maps of NorA-BRIL-BAG2-I-23 (using 302,069 final particles) as assessed using the gold standard Fourier shell correlation (FSC) at FSC = 0.143 ^61^. Particles were evenly distributed with various orientations that generated 2D classes of similar resolution. Final maps were masked around NorA-I-23 and locally refined to obtain a final 2.79 Å resolution map.

### Structural model refinement

The initial model of the complex was generated using ModelAngelo ^62^ and the final model was then refined against both non-uniform and locally refined maps using Coot ^63^ and Phenix ^64^.

## Acknowledgements

This work was supported by NIH awards (R01AI108889, R01AI165782, R21AI188334) to N.J.T, Defense Advanced Research Projects Agency (HR0011-21-2-0012) and Open Philanthropy Project Improving Protein Design Fund grants to A.C.G, and the Howard Hughes Medical Institute to D.B. We thank David Hooper for sharing the pTrcHis2C vector encoding *norA*, Rico Rojas and Tahj Starr for recombineering discussions in *E. coli*, Tiffany Suwatthee for performing preliminary growth inhibition experiments in *S. aureus*, Huihui Kuang and William Rice for assistance at the NYU Langone Health Cryo-Electron Microscopy Laboratory, and Shenglong Wang for assistance at the NYU High Performance Computing (HPC) facility. Cryo-EM data were collected at NYU Langone Health’s Cryo-Electron Microscopy Laboratory (RRID: SCR_019202), which is partially supported by the Laura and Isaac Perlmutter Cancer Center Support Grant NIH/NCI P30CA016087 as well as NIH Grant R01NS108151. Cryo-EM data were processed using the NYU HPC facility. The pBAD33-Gm vector (plasmid #65098; http://n2t.net/addgene:65098; RRID: Addgene_65098) and pSIJ8 vector (plasmid # 68122; http://n2t.net/addgene:68122; RRID: Addgene_68122) were obtained through Addgene.

## Author Contributions

P.M. created the E. coli Δ*acrB* mutant, optimized the *E. coli* co-expression experiment, performed and quantified the co-expression screen, performed pull down experiments and immunoblot analyses, created alanine mutants of I-23 and performed the functional assays on I-23, prepared the cryo-EM sample, and wrote the manuscript. A.C.G. conceived of the idea to use miniproteins and computational protein design to inhibit NorA, designed the miniprotein library, performed computation/experiments to improve miniprotein solubility, and helped prepare the manuscript. G.L. prepared the cryo-EM sample, processed and analyzed cryo-EM datasets, and determined the cryo-EM structure. D.K. created the Autograft loop grafting script used in the miniprotein design. D.B. directed the project. N.J.T. designed and directed the project and wrote the manuscript. All authors participated in data analysis and revision of the manuscript.

## Data Availability

The cryo-EM map has been deposited in the Electron Microscopy Data Bank (EMDB) under accession code EMD-56885 (NorA in the outward-open conformation bound to miniprotein I-23). The atomic coordinates have been deposited in the Protein Data Bank (PDB) under accession code 28VJ (NorA in the outward-open conformation bound to miniprotein I-23).

## Supporting Information

Figures include miniprotein screening results, pull down experiments, purification of the NorA-BRIL-BAG2-I-23 complex for cryo-EM, data processing workflow for cryo-EM, details on the cryo-EM data collection and results, structural overlays, and the *S. aureus* growth inhibition results. Tables include *IC_50_* values for the miniproteins, sequences for the top 10 miniproteins, and cryo-EM structural refinement statistics.

## References

(1) Turner, NA, Sharma-Kuinkel, BK, Maskarinec, SA, Eichenberger, EM, Shah, PP, Carugati, M, Holland, TL, Fowler, VG, Jr. Methicillin-resistant Staphylococcus aureus: an overview of basic and clinical research. Nature reviews. Microbiology 2019, 17 (4), 203–218. DOI: 10.1038/s41579-018-0147-4 From NLM Medline.

(2) Hiramatsu, K, Katayama, Y, Matsuo, M, Sasaki, T, Morimoto, Y, Sekiguchi, A, Baba, T. Multi-drug-resistant Staphylococcus aureus and future chemotherapy. Journal of Infection and Chemotherapy 2014, 20 (10), 593–601. DOI: 10.1016/j.jiac.2014.08.001 (acccessed 2025/08/20/15:44:25). From DOI.org (Crossref).

(3) Scully, J, Mustafa, AS, Hanif, A, Tunio, JH, Tunio, SNJ. Immune Responses to Methicillin-Resistant Staphylococcus aureus Infections and Advances in the Development of Vaccines and Immunotherapies. Vaccines 2024, 12 (10), 1106. DOI: 10.3390/vaccines12101106 (acccessed 2025/08/20/15:45:36). From DOI.org (Crossref).

(4) Barnabas, V, Kashyap, A, Raja, R, Newar, K, Rai, D, Dixit, NM, Mehra, S. The Extent of Antimicrobial Resistance Due to Efflux Pump Regulation. ACS Infect Dis 2022, 8 (11), 2374–2388. DOI: 10.1021/acsinfecdis.2c00460.

(5) Kapoor, G, Saigal, S, Elongavan, A. Action and resistance mechanisms of antibiotics: A guide for clinicians. J Anaesthesiol Clin Pharmacol 2017, 33 (3), 300–305. DOI: 10.4103/joacp.JOACP_349_15.

(6) Kumawat, M, Nabi, B, Daswani, M, Viquar, I, Pal, N, Sharma, P, Tiwari, S, Sarma, DK, Shubham, S, Kumar, M. Role of bacterial efflux pump proteins in antibiotic resistance across microbial species. Microb Pathog 2023, 181, 106182. DOI: 10.1016/j.micpath.2023.106182.

(7) Zhang, L, Tian, X, Sun, L, Mi, K, Wang, R, Gong, F, Huang, L. Bacterial Efflux Pump Inhibitors Reduce Antibiotic Resistance. Pharmaceutics 2024, 16 (2). DOI: 10.3390/pharmaceutics16020170.

(8) Brawley, DN, Sauer, DB, Li, J, Zheng, X, Koide, A, Jedhe, GS, Suwatthee, T, Song, J, Liu, Z, Arora, PS, Koide, S, Torres, VJ, Wang, DN, Traaseth, NJ. Structural basis for inhibition of the drug efflux pump NorA from Staphylococcus aureus. Nat Chem Biol 2022, 18 (7), 706–712. DOI: 10.1038/s41589-022-00994-9.

(9) Mitchell, P. A general theory of membrane transport from studies of bacteria. Nature 1957, 180 (4577), 134–136. DOI: 10.1038/180134a0.

(10) Latorraca, NR, Fastman, NM, Venkatakrishnan, AJ, Frommer, WB, Dror, RO, Feng, L. Mechanism of Substrate Translocation in an Alternating Access Transporter. Cell 2017, 169 (1), 96–107. DOI: 10.1016/j.cell.2017.03.010.

(11) Jardetzky, O. Simple allosteric model for membrane pumps. Nature 1966, 211 (5052), 969–970.

(12) Kathawala, RJ, Gupta, P, Ashby, CR, Jr., Chen, ZS. The modulation of ABC transporter-mediated multidrug resistance in cancer: a review of the past decade. Drug Resist Updat 2015, 18, 1–17. DOI: 10.1016/j.drup.2014.11.002.

(13) Renau, TE, Leger, R, Filonova, L, Flamme, EM, Wang, M, Yen, R, Madsen, D, Griffith, D, Chamberland, S, Dudley, MN;, et al. Conformationally-restricted analogues of efflux pump inhibitors that potentiate the activity of levofloxacin in *Pseudomonas aeruginosa*. Bioorg. Med. Chem. Lett. 2003, 13 (16), 2755–2758. DOI: 10.1016/S0960-894x(03)00556-0.

(14) Lomovskaya, O, Bostian, KA. Practical applications and feasibility of efflux pump inhibitors in the clinic--a vision for applied use. Biochem Pharmacol 2006, 71 (7), 910–918. DOI: 10.1016/j.bcp.2005.12.008 From NLM Medline.

(15) Kim, J, Tan, YZ, Wicht, KJ, Erramilli, SK, Dhingra, SK, Okombo, J, Vendome, J, Hagenah, LM, Giacometti, SI, Warren, AL;, et al. Structure and drug resistance of the *Plasmodium falciparum* transporter PfCRT. Nature 2019, 576 (7786), 315–320. DOI: 10.1038/s41586-019-1795-x.

(16) Debruycker, V, Hutchin, A, Masureel, M, Ficici, E, Martens, C, Legrand, P, Stein, RA, McHaourab, HS, Faraldo-Gomez, JD, Remaut, H, Govaerts, C. An embedded lipid in the multidrug transporter LmrP suggests a mechanism for polyspecificity. Nat Struct Mol Biol 2020, 27 (9), 829–835. DOI: 10.1038/s41594-020-0464-y.

(17) Meng, X, Li, T, Zhao, Y, Wu, C. CXC-Mediated Cellular Uptake of Miniproteins: Forsaking “Arginine Magic”. ACS Chem. Biol. 2018, 13 (11), 3078–3086. DOI: 10.1021/acschembio.8b00564 (acccessed 2025/08/22/16:14:33). From DOI.org (Crossref).

(18) Ciesiołkiewicz, A, Lizandra Perez, J, Berlicki, Ł. Miniproteins in medicinal chemistry. Bioorganic & Medicinal Chemistry Letters 2022, 71, 128806. DOI: 10.1016/j.bmcl.2022.128806 (acccessed 2025/08/22/15:43:46). From DOI.org (Crossref).

(19) Crook, ZR, Nairn, NW, Olson, JM. Miniproteins as a Powerful Modality in Drug Development. Trends in Biochemical Sciences 2020, 45 (4), 332–346. DOI: 10.1016/j.tibs.2019.12.008 (acccessed 2025/08/22/15:45:32). From DOI.org (Crossref).

(20) Daisy Precilla, S, Biswas, I, Anitha, TS, Agieshkumar, B. Microproteins unveiling new dimensions in cancer. Funct Integr Genomics 2024, 24 (5), 152. DOI: 10.1007/s10142-024-01426-8 (acccessed 2025/08/22/16:28:03). From DOI.org (Crossref).

(21) Colom, MS, Michelet, X, Bahl, CD. 1439 De novo design and optimization of multivalent miniprotein NK cell engagers targeting acute myeloid leukemia. In SITC 39th Annual Meeting (SITC 2024) Abstracts, 2024/11//, 2024; BMJ Publishing Group Ltd: pp A1610-A1610. DOI: 10.1136/jitc-2024-SITC2024.1439.

(22) Asada, N, Krebs, CF, Panzer, U. Miniproteins may have a big impact: new therapeutics for autoimmune diseases and beyond. Sig Transduct Target Ther 2024, 9 (1), 298. DOI: 10.1038/s41392-024-02010-z (acccessed 2025/08/22/15:40:20). From DOI.org (Crossref).

(23) Ożga, K, Berlicki, Ł. Design and Engineering of Miniproteins. ACS Bio Med Chem Au 2022, 2 (4), 316–327. DOI: 10.1021/acsbiomedchemau.2c00008 (acccessed 2025/08/22/16:12:01). From DOI.org (Crossref).

(24) Case, JB, Chen, RE, Cao, L, Ying, B, Winkler, ES, Johnson, M, Goreshnik, I, Pham, MN, Shrihari, S, Kafai, NM;, et al. Ultrapotent miniproteins targeting the SARS-CoV-2 receptor-binding domain protect against infection and disease. Cell Host & Microbe 2021, 29 (7), 1151–1161.e1155. DOI: 10.1016/j.chom.2021.06.008 (acccessed 2025/08/20/15:51:47). From DOI.org (Crossref).

(25) Ragotte, RJ, Tortorici, MA, Catanzaro, NJ, Addetia, A, Coventry, B, Froggatt, HM, Lee, J, Stewart, C, Brown, JT, Goreshnik, I;, et al. Designed miniproteins potently inhibit and protect against MERS-CoV. Cell Reports 2025, 44 (6), 115760. DOI: 10.1016/j.celrep.2025.115760 (acccessed 2025/08/20/15:54:21). From DOI.org (Crossref).

(26) Berger, S, Seeger, F, Yu, T-Y, Aydin, M, Yang, H, Rosenblum, D, Guenin-Macé, L, Glassman, C, Arguinchona, L, Sniezek, C;, et al. Preclinical proof of principle for orally delivered Th17 antagonist miniproteins. Cell 2024, 187 (16), 4305–4317.e4318. DOI: 10.1016/j.cell.2024.05.052 (acccessed 2025/08/20/15:50:45). From DOI.org (Crossref).

(27) Crunkhorn, S. Oral miniproteins treat IBD. Nat Rev Drug Discov 2024, 23 (9), 660–660. DOI: 10.1038/d41573-024-00126-z (acccessed 2025/08/22/16:24:31). From DOI.org (Crossref).

(28) Yu, JL, Grinius, L, Hooper, DC. NorA functions as a multidrug efflux protein in both cytoplasmic membrane vesicles and reconstituted proteoliposomes. Journal of bacteriology 2002, 184 (5), 1370–1377, Research Support, U.S. Gov’t, P.H.S.

(29) Neyfakh, AA, Borsch, CM, Kaatz, GW. Fluoroquinolone resistance protein NorA of *Staphylococcus aureus* is a multidrug efflux transporter. Antimicrobial agents and chemotherapy 1993, 37 (1), 128–129, Research Support, Non-U.S. Gov’t.

(30) Yoshida, H, Bogaki, M, Nakamura, S, Ubukata, K, Konno, M. Nucleotide sequence and characterization of the *Staphylococcus aureus* norA gene, which confers resistance to quinolones. Journal of bacteriology 1990, 172 (12), 6942–6949. DOI: 10.1128/jb.172.12.6942-6949.1990.

(31) Yu, XH, Hao, ZH, Liu, PL, Liu, MM, Zhao, LL, Zhao, X. Increased Expression of Efflux Pump norA Drives the Rapid Evolutionary Trajectory from Tolerance to Resistance against Ciprofloxacin in Staphylococcus aureus. Antimicrobial agents and chemotherapy 2022, 66 (12), e0059422. DOI: 10.1128/aac.00594-22.

(32) Tintino, SR, Wilairatana, P, De Souza, VCA, Da Silva, JMA, Pereira, PS, De Morais Oliveira-Tintino, CD, De Matos, YMLS, Júnior, JTC, De Queiroz Balbino, V, Siqueira-Junior, JP, Menezes, IRA, Siyadatpanah, A, Coutinho, HDM, Balbino, TCL. Inhibition of the norA gene expression and the NorA efflux pump by the tannic acid. Sci Rep 2023, 13 (1), 17394. DOI: 10.1038/s41598-023-43038-5 (acccessed 2025/08/20/15:25:14). From DOI.org (Crossref).

(33) Ng, EY, Trucksis, M, Hooper, DC. Quinolone resistance mediated by norA: physiologic characterization and relationship to flqB, a quinolone resistance locus on the Staphylococcus aureus chromosome. Antimicrob Agents Chemother 1994, 38 (6), 1345–1355. DOI: 10.1128/AAC.38.6.1345 (acccessed 2025/08/20/15:46:19). From DOI.org (Crossref).

(34) Pasquina-Lemonche, L, Burns, J, Turner, RD, Kumar, S, Tank, R, Mullin, N, Wilson, JS, Chakrabarti, B, Bullough, PA, Foster, SJ, Hobbs, JK. The architecture of the Gram-positive bacterial cell wall. Nature 2020, 582 (7811), 294–297. DOI: 10.1038/s41586-020-2236-6 From NLM Medline.

(35) Watson, JL, Juergens, D, Bennett, NR, Trippe, BL, Yim, J, Eisenach, HE, Ahern, W, Borst, AJ, Ragotte, RJ, Milles, LF;, et al. De novo design of protein structure and function with RFdiffusion. Nature 2023, 620 (7976), 1089–1100. DOI: 10.1038/s41586-023-06415-8 From NLM Medline.

(36) Dauparas, J, Anishchenko, I, Bennett, N, Bai, H, Ragotte, RJ, Milles, LF, Wicky, BIM, Courbet, A, de Haas, RJ, Bethel, N;, et al. Robust deep learning-based protein sequence design using ProteinMPNN. Science 2022, 378 (6615), 49–56. DOI: 10.1126/science.add2187 From NLM Medline.

(37) Guo, HB, Perminov, A, Bekele, S, Kedziora, G, Farajollahi, S, Varaljay, V, Hinkle, K, Molinero, V, Meister, K, Hung, C, Dennis, P, Kelley-Loughnane, N, Berry, R. AlphaFold2 models indicate that protein sequence determines both structure and dynamics. Scientific reports 2022, 12 (1), 10696. DOI: 10.1038/s41598-022-14382-9.

(38) Bennett, NR, Coventry, B, Goreshnik, I, Huang, B, Allen, A, Vafeados, D, Peng, YP, Dauparas, J, Baek, M, Stewart, L, DiMaio, F, De Munck, S, Savvides, SN, Baker, D. Improving de novo protein binder design with deep learning. Nat Commun 2023, 14 (1), 2625. DOI: 10.1038/s41467-023-38328-5 From NLM Medline.

(39) Li, J, Li, Y, Koide, A, Kuang, H, Torres, VJ, Koide, S, Wang, DN, Traaseth, NJ. Proton-coupled transport mechanism of the efflux pump NorA. Nat Commun 2024, 15 (1), 4494. DOI: 10.1038/s41467-024-48759-3.

(40) Ethier, M, Lambert, J-P, Vasilescu, J, Figeys, D. Analysis of protein interaction networks using mass spectrometry compatible techniques. Analytica Chimica Acta 2006, 564 (1), 10–18. DOI: 10.1016/j.aca.2005.12.046 (acccessed 2025/08/20/22:17:30). From DOI.org (Crossref).

(41) Valdez-Sinon, AN, Gokhale, A, Faundez, V, Bassell, GJ. Protocol for Immuno-Enrichment of FLAG-Tagged Protein Complexes. STAR Protocols 2020, 1 (2), 100083. DOI: 10.1016/j.xpro.2020.100083 (acccessed 2025/08/20/22:22:22). From DOI.org (Crossref).

(42) Gerace, E, Moazed, D. Affinity Pull-Down of Proteins Using Anti-FLAG M2 Agarose Beads. In Methods in Enzymology, Vol. 559; Elsevier, 2015; pp 99–110.

(43) Zhao, X, Li, G, Liang, S. Several Affinity Tags Commonly Used in Chromatographic Purification. Journal of Analytical Methods in Chemistry 2013, 2013, 1–8. DOI: 10.1155/2013/581093 (acccessed 2025/08/20/22:15:34). From DOI.org (Crossref).

(44) Xie, P, Li, Y, Lamon, G, Kuang, H, Wang, DN, Traaseth, NJ. A fiducial-assisted strategy compatible with resolving small MFS transporter structures in multiple conformations using cryo-EM. Nat Commun 2025, 16 (1), 7. DOI: 10.1038/s41467-024-54986-5 From NLM Medline.

(45) Mukherjee, S, Erramilli, SK, Ammirati, M, Alvarez, FJD, Fennell, KF, Purdy, MD, Skrobek, BM, Radziwon, K, Coukos, J, Kang, Y;, et al. Synthetic antibodies against BRIL as universal fiducial marks for single-particle cryoEM structure determination of membrane proteins. Nat Commun 2020, 11 (1), 1598. DOI: 10.1038/s41467-020-15363-0.

(46) Lynn C Thomason, NC, Xintian Li, Donald L Court. Recombineering: Genetic Engineering in E. coli Using Homologous Recombination. Curr Protoc. 2024. DOI: 10.1002/cpz1.656.

(47) Sheila I. Jensen, RML, Markus J. Herrgård & Alex T. Nielsen Seven gene deletions in seven days: Fast generation of Escherichia coli strains tolerant to acetate and osmotic stress. Sci Rep. DOI: 10.1038/srep17874.

(48) K A Datsenko 1, BLW. One-step inactivation of chromosomal genes in Escherichia coli K-12 using PCR products. PNAS 2000. DOI: 10.1073/pnas.120163297.

(49) Jimenez, N, Lacasta, A, Vilches, S, Reyes, M, Vazquez, J, Aquillini, E, Merino, S, Regue, M, Tomas, JM. Genetics and proteomics of *Aeromonas salmonicida* lipopolysaccharide core biosynthesis. Journal of bacteriology 2009, 191 (7), 2228–2236. DOI: 10.1128/JB.01395-08.

(50) Bailey, LJ, Sheehy, KM, Dominik, PK, Liang, WG, Rui, H, Clark, M, Jaskolowski, M, Kim, Y, Deneka, D, Tang, W-J, Kossiakoff, AA. Locking the Elbow: Improved Antibody Fab Fragments as Chaperones for Structure Determination. Journal of Molecular Biology 2018, 430 (3), 337–347. DOI: 10.1016/j.jmb.2017.12.012 (acccessed 2025/08/21/17:59:19). From DOI.org (Crossref).

(51) Rheinberger, J, Oostergetel, G, Resch, GP, Paulino, C. Optimized cryo-EM data-acquisition workflow by sample-thickness determination. Acta Crystallogr D Struct Biol 2021, 77 (5), 565–571. DOI: 10.1107/S205979832100334X (acccessed 2025/08/21/18:01:34). From DOI.org (Crossref).

(52) Rice, WJ, Cheng, A, Noble, AJ, Eng, ET, Kim, LY, Carragher, B, Potter, CS. Routine determination of ice thickness for cryo-EM grids. Journal of Structural Biology 2018, 204 (1), 38–44. DOI: 10.1016/j.jsb.2018.06.007 (acccessed 2025/08/21/18:03:08). From DOI.org (Crossref).

(53) Suloway, C, Pulokas, J, Fellmann, D, Cheng, A, Guerra, F, Quispe, J, Stagg, S, Potter, CS, Carragher, B. Automated molecular microscopy: The new Leginon system. Journal of Structural Biology 2005, 151 (1), 41–60. DOI: 10.1016/j.jsb.2005.03.010 (acccessed 2025/08/21/18:02:24). From DOI.org (Crossref).

(54) Cheng, A, Negro, C, Bruhn, JF, Rice, WJ, Dallakyan, S, Eng, ET, Waterman, DG, Potter, CS, Carragher, B. Leginon: New features and applications. Protein Science 2021, 30 (1), 136–150. DOI: 10.1002/pro.3967 (acccessed 2025/08/21/18:05:05). From DOI.org (Crossref).

(55) Rohou, A, Grigorieff, N. CTFFIND4: Fast and accurate defocus estimation from electron micrographs. Journal of Structural Biology 2015, 192 (2), 216–221. DOI: 10.1016/j.jsb.2015.08.008 (acccessed 2025/08/21/18:08:35). From DOI.org (Crossref).

(56) Zheng, SQ, Palovcak, E, Armache, J-P, Verba, KA, Cheng, Y, Agard, DA. MotionCor2: anisotropic correction of beam-induced motion for improved cryo-electron microscopy. Nat Methods 2017, 14 (4), 331–332. DOI: 10.1038/nmeth.4193 (acccessed 2025/08/21/18:07:20). From DOI.org (Crossref).

(57) Lander, GC, Stagg, SM, Voss, NR, Cheng, A, Fellmann, D, Pulokas, J, Yoshioka, C, Irving, C, Mulder, A, Lau, P-W, Lyumkis, D, Potter, CS, Carragher, B. Appion: An integrated, database-driven pipeline to facilitate EM image processing. Journal of Structural Biology 2009, 166 (1), 95–102. DOI: 10.1016/j.jsb.2009.01.002 (acccessed 2025/08/21/18:15:07). From DOI.org (Crossref).

(58) Tegunov, D, Cramer, P. Real-time cryo-electron microscopy data preprocessing with Warp. Nat Methods 2019, 16 (11), 1146–1152. DOI: 10.1038/s41592-019-0580-y (accessed 2025/08/21/18:16:09). From DOI.org (Crossref).

(59) Punjani, A, Rubinstein, JL, Fleet, DJ, Brubaker, MA. cryoSPARC: algorithms for rapid unsupervised cryo-EM structure determination. Nat Methods 2017, 14 (3), 290–296. DOI: 10.1038/nmeth.4169 (acccessed 2025/08/21/18:19:56). From DOI.org (Crossref).

(60) Bepler, T, Morin, A, Rapp, M, Brasch, J, Shapiro, L, Noble, AJ, Berger, B. Positive-unlabeled convolutional neural networks for particle picking in cryo-electron micrographs. Nat Methods 2019, 16 (11), 1153–1160. DOI: 10.1038/s41592-019-0575-8 (acccessed 2025/08/21/18:18:05). From DOI.org (Crossref).

(61) Rosenthal, PB, Henderson, R. Optimal Determination of Particle Orientation, Absolute Hand, and Contrast Loss in Single-particle Electron Cryomicroscopy. Journal of Molecular Biology 2003, 333 (4), 721–745. DOI: 10.1016/j.jmb.2003.07.013 (acccessed 2025/08/21/18:23:37). From DOI.org (Crossref).

(62) Jamali, K, Kall, L, Zhang, R, Brown, A, Kimanius, D, Scheres, SHW. Automated model building and protein identification in cryo-EM maps. Nature 2024, 628 (8007), 450–457. DOI: 10.1038/s41586-024-07215-4 From NLM Medline.

(63) Emsley, P, Cowtan, K. Coot: model-building tools for molecular graphics. Acta Crystallogr D Biol Crystallogr 2004, 60 (Pt 12 Pt 1), 2126-2132. DOI: 10.1107/S0907444904019158.

(64) Adams, PD, Afonine, PV, Bunkoczi, G, Chen, VB, Davis, IW, Echols, N, Headd, JJ, Hung, LW, Kapral, GJ, Grosse-Kunstleve, RW;, et al. PHENIX: a comprehensive Python-based system for macromolecular structure solution. Acta Crystallogr D Biol Crystallogr 2010, 66 (Pt 2), 213–221. DOI: 10.1107/S0907444909052925 From NLM Medline.

