## Supporting Information for "Miniprotein inhibitors of the *Staphylococcus aureus* efflux transporter NorA"

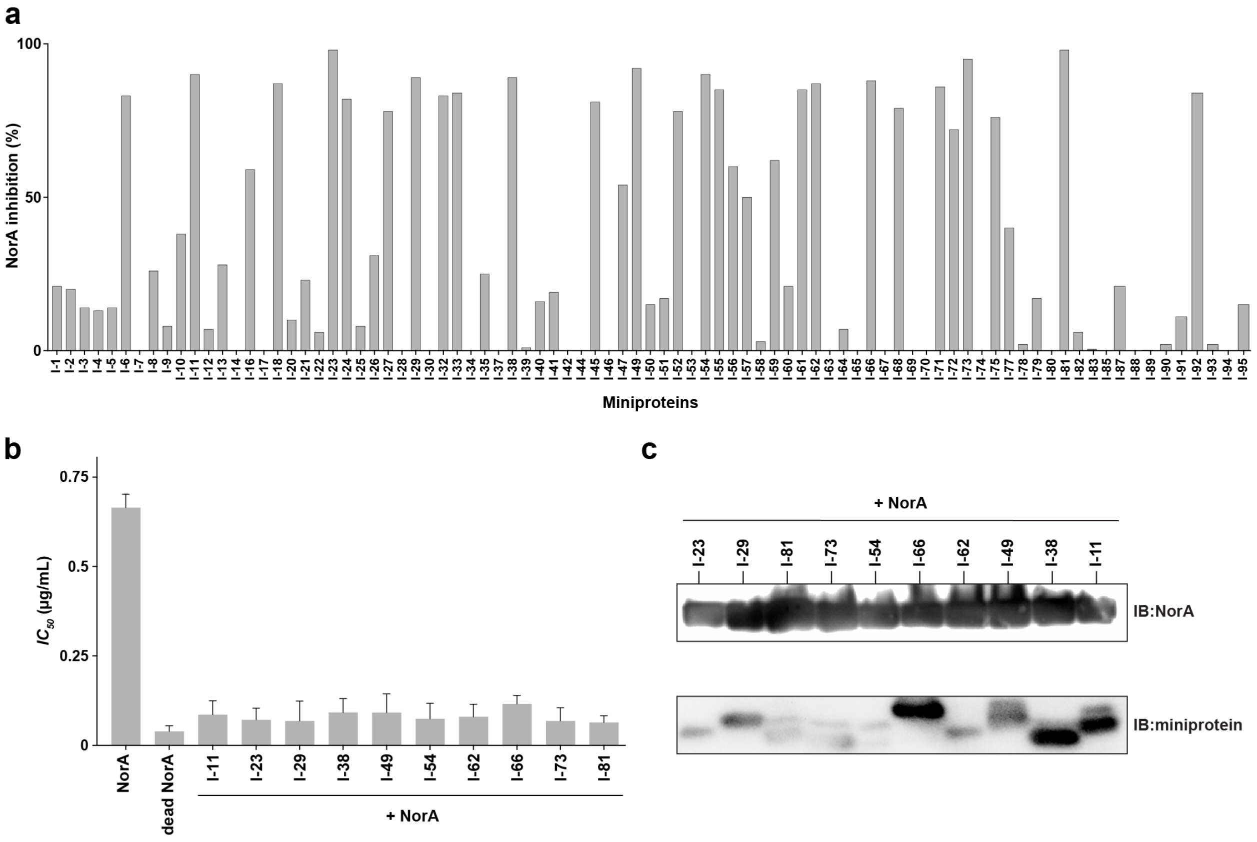


**Figure S1. Screening results and co-immunoprecipitation of miniproteins following NorA purification.**

**a.** *E. coli* co-expression growth inhibition screening results for the 86-member miniprotein library. The $IC_{50}$ values for norfloxacin were converted into a % NorA inhibition using the NorA and dead NorA (E222A) expressed samples as references.

**b.** Growth inhibition $IC_{50}$ values of norfloxacin for NorA co-expressed with FLAG-tagged miniproteins. No change in inhibition was observed following incorporation of the FLAG-tag at the miniprotein C-terminus.

**c.** Immunoblotting (IB) analyses performed for NorA (C-terminal MYC-tag) and miniproteins (C-terminal FLAG-tag) following NorA purification (i.e., isolating the membrane fraction, solubilization in LMNG detergent, and passing over a Ni-NTA column).

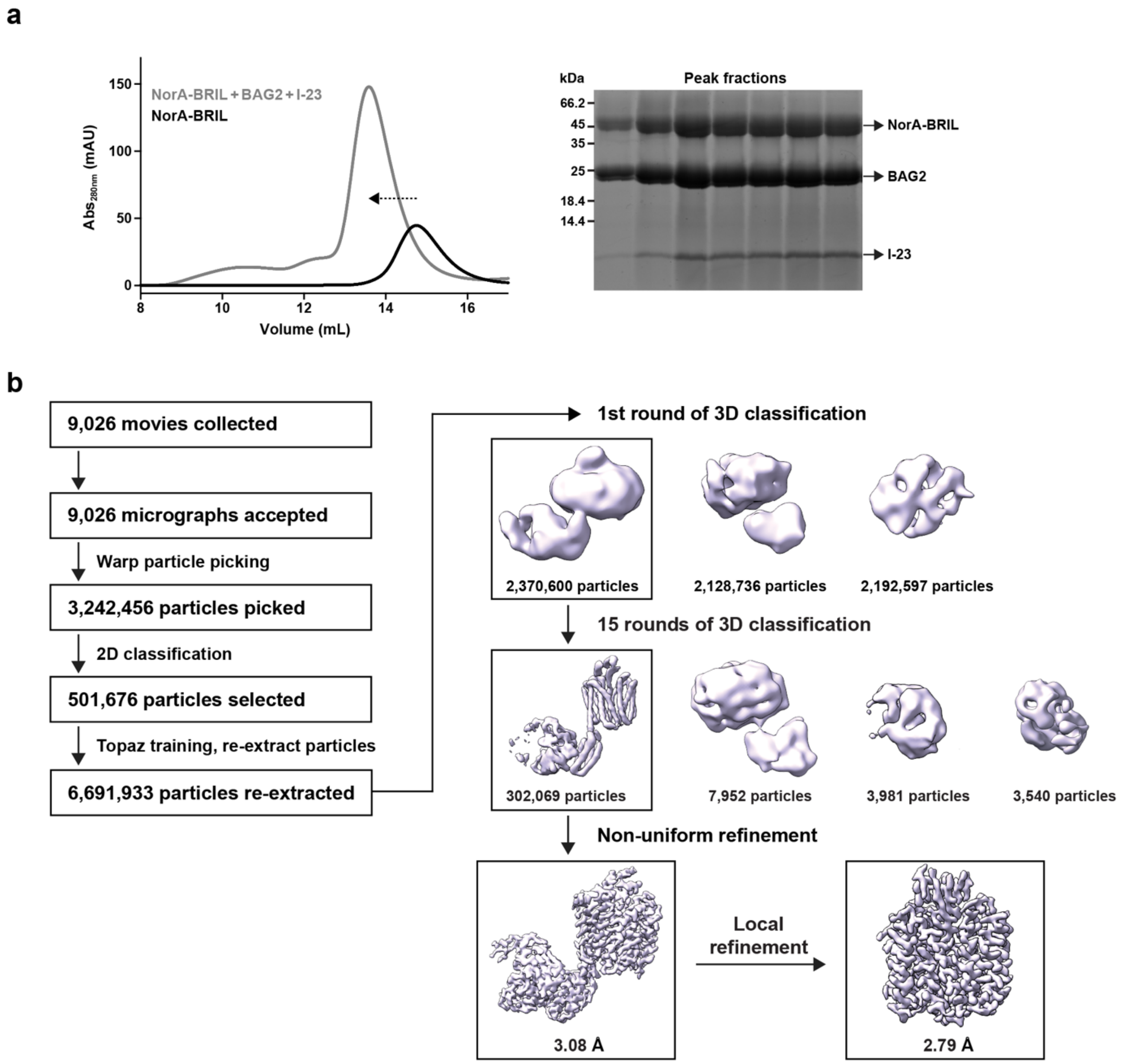


**Figure S2. Sample preparation and cryo-EM processing of the NorA-BRIL-BAG2-I-23 complex.**

**a.** Left: SEC chromatogram of NorA-BRIL (black) and NorA-BRIL in the presence of BAG2 and I-23 (grey). The left shifted peak is indicated by the arrow. Right: SDS-PAGE gel of the peak fractions corresponding to the SEC peak for the NorA-BRIL-BAG2-I-23 complex.

**b.** Cryo-EM processing pipeline of the dataset collected on the NorA-BRIL-BAG2-I-23 complex. Note that an additional 4$^{th}$ "junk" class was added in the middle of heterogeneous refinement.

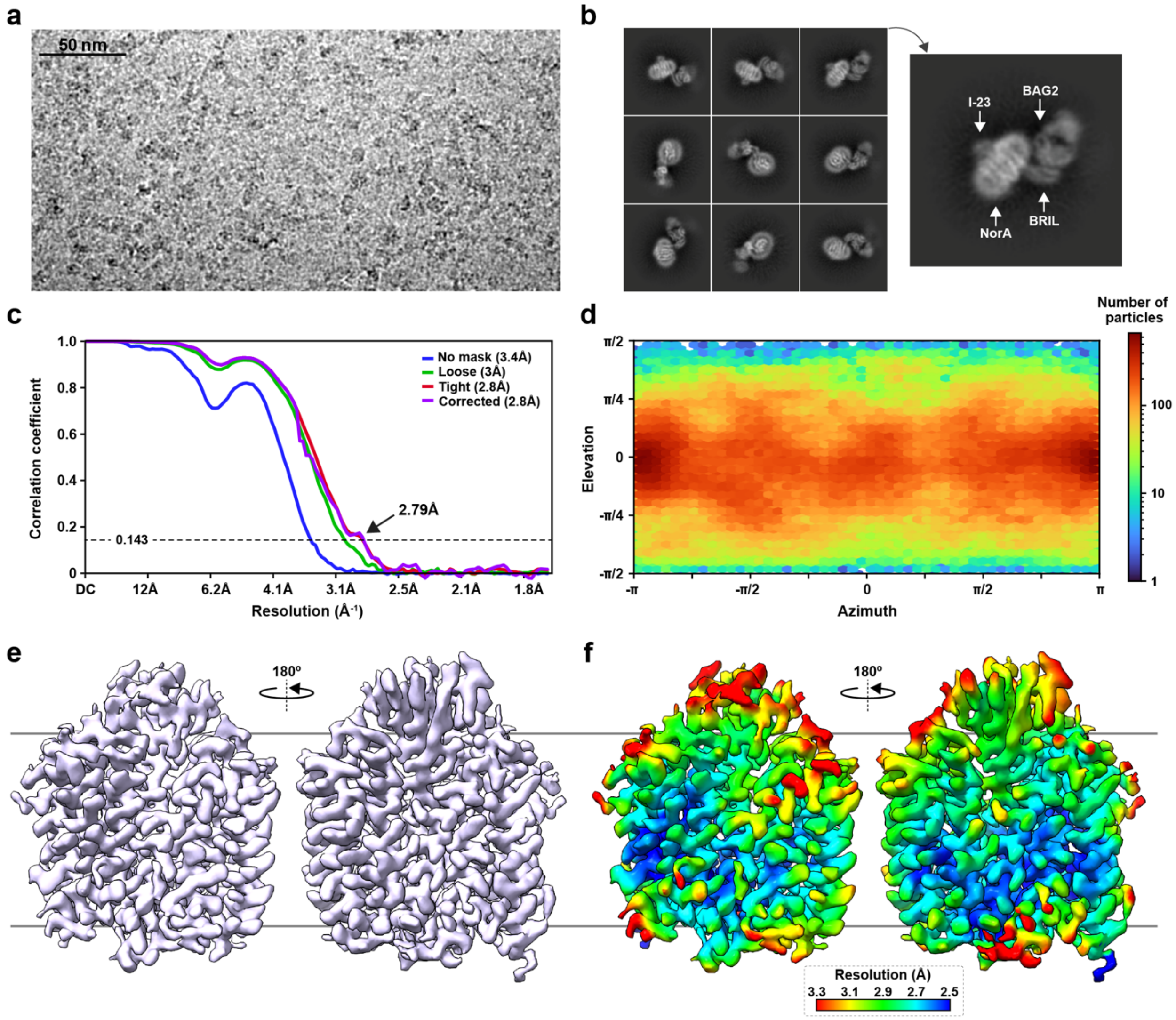


**Figure S3. Cryo-EM structure determination of the NorA-I-23 complex.**

**a.** Representative micrograph from the cryo-EM dataset.

**b.** Exemplary 2D classes from the final particle stack used for 3D reconstruction. An annotated class displays the locations of NorA, BRIL, BAG2, and I-23.

**c.** Fourier shell correlation curves corresponding to the NorA-I-23 reconstruction.

**d.** Orientation distribution heatmaps for the NorA-I-23 reconstruction.

**e, f.** Side views of the Coulomb potential map (e) and local resolution map (f) of the NorA-I-23 map. The local resolution map is displayed using a linear coloring scale.

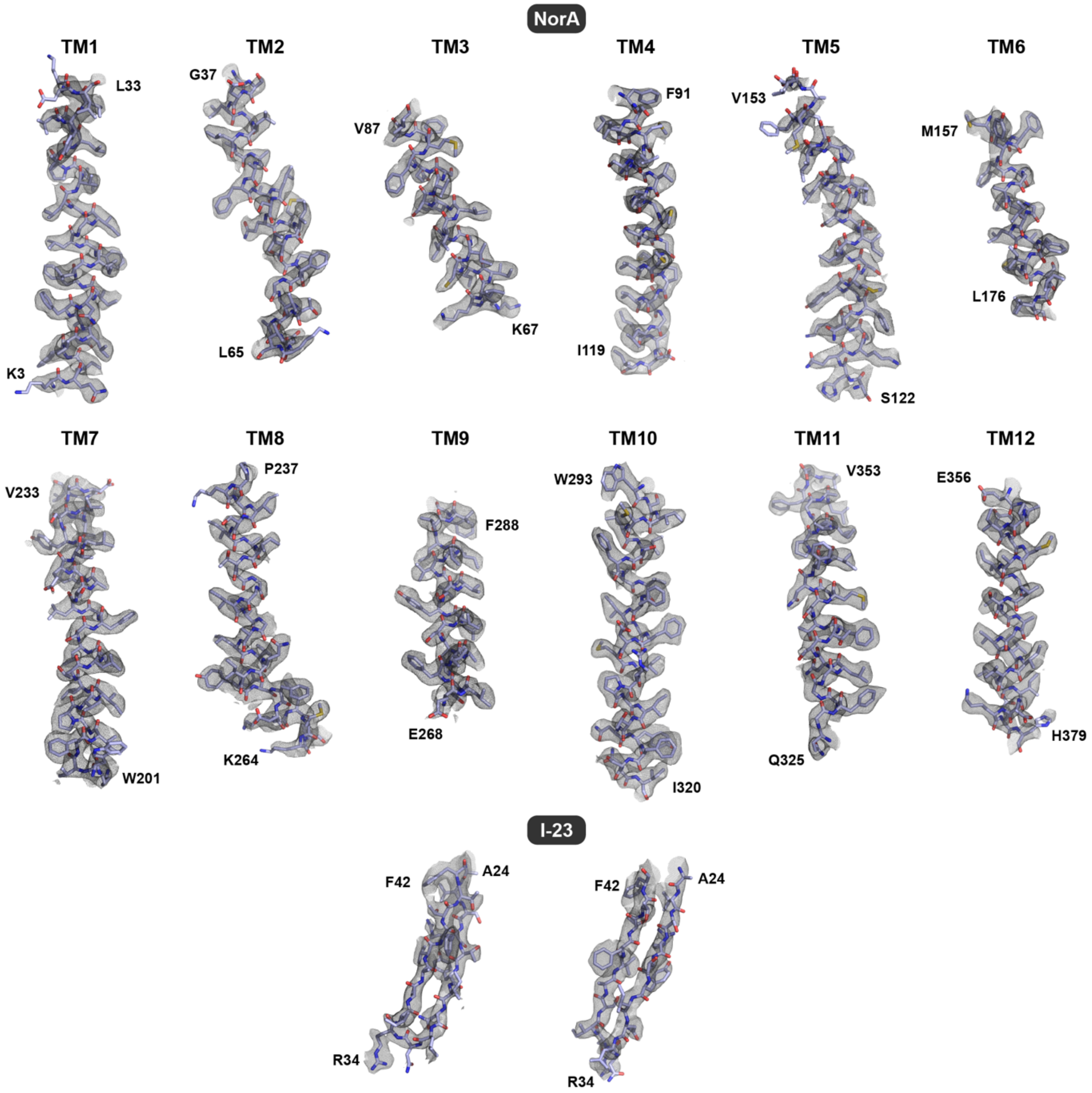


**Figure S4. Quality assessment of the NorA-I-23 structural model.**

Model-to-map fitting displaying superimpositions of the cryo-EM map (grey mesh) and the structural model (light blue stick representation). NorA TM helices are displayed in the top and middle rows while the I-23 β-hairpin loop is displayed in the bottom row with two different views. The map contour level was set to 10σ using the isomesh command in PyMOL. The indicated residues define the segments displayed.

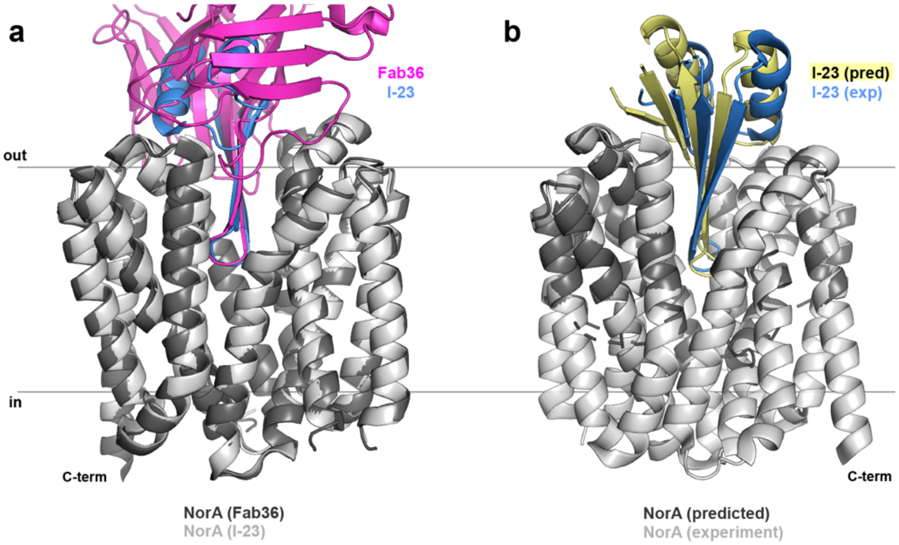


**Figure S5. Structural comparison of NorA-I-23 with the NorA-Fab36 structure and predicted model.**

**a.** Superimposition of NorA-Fab36 (PDB ID: 9B3K) and NorA-I-23 (*this work*) aligned using NorA from each model. The backbone RMSD is 0.459 Å. Fab36 is displayed in magenta, I-23 in light blue, and NorA is displayed in dark (in the NorA-Fab36 complex) or light grey (in the NorA-I-23 complex).

**b.** Superimposition of experimental and predicted NorA-I-23 complex structures. The backbone RMSD is 0.681 Å. Experimental I-23 is in light blue, predicted I-23 is in yellow, experimental NorA is in light grey, and predicted NorA is in dark grey. Note that predicted NorA contains only a portion of the entire structure to expedite miniprotein design calculations.

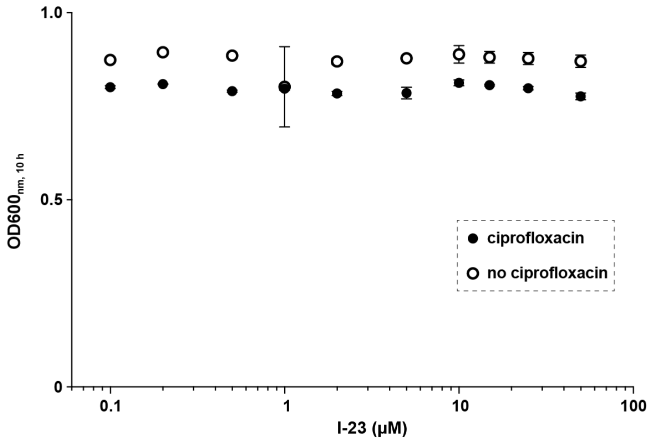


**Figure S6.** ***S. aureus*** **growth inhibition results in the presence of I-23 and ciprofloxacin.** Growth inhibition experiments of *S. aureus* SA1199B at 10 h in the presence (black circles) or absence (white circles) of 1 µg/mL ciprofloxacin and varying concentrations of I-23. No growth inhibition in the presence of ciprofloxacin was observed at the highest concentration of I-23 tested (50 µM). Experiments were repeated in two independent experiments with error bars showing the standard deviation among replicate data points.

**Table S1. Miniprotein inhibition of NorA displayed in the norfloxacin $IC_{50}$ value obtained from growth inhibition experiments. Errors reflect values obtained in duplicate.**

| Sample | $IC_{50}$ (µg/mL) |
|---|---|
| NorA | 0.7442 ± 0.061 |
| $NorA^{E222A}$ | 0.0405 ± 0.013 |
| NorA + I-81 | 0.059 ± 0.01 |
| NorA + I-23 | 0.062 ± 0.017 |
| NorA + I-73 | 0.105 ± 0.009 |
| NorA + I-49 | 0.098 ± 0.012 |
| NorA + I-54 | 0.101 ± 0.010 |
| NorA + I-11 | 0.105 ± 0.012 |
| NorA + I-62 | 0.115 ± 0.012 |
| NorA + I-29 | 0.129 ± 0.009 |
| NorA + I-38 | 0.134 ± 0.004 |
| NorA + I-66 | 0.1534 ± 0.0006 |

**Table S2. Protein sequences of the top 10 miniprotein hits from the screen. Underlined portions indicate a shared sequence with the CDRH3 of Fab36.**

| Miniprotein | Protein sequence |
|---|---|
| I-11 | PTIIAEVRSLSKRKKLIELAKELGAKTTEVFYYAWRVGGVYLSVETEDE<br>EKADKIEEKAKELGLKVWVSR |
| I-23 | MYEVIVRNAPRSFVKEVREETGAKVSRTYINLNRISAVFTTFTHERKED<br>AEAIAERARARGLEVFLVE |
| I-29 | SMLEEENKLYEEYEKARKELIAKMKELGAKRLYYYAWRVGGIYLVTP<br>DKVAGISYGVINKAFARYQLRVLKETLEEL |
| I-38 | TEIILTGISLRKALKFAKEEGCKVSFTFYYAWRVGGVYVTLTGIDEEKAR<br>KFAEEEGVEVYVIK |
| I-49 | MQVIIANANAKVLKAAKELAKEKGVKLTRTYYYAWRVGGIFITLEGVS<br>KEDAEELTALAKEEGCIVEVIE |
| I-54 | MYQVIVVNATKELIRYAKSIPGVKIERVYFPAFRASAIYTVFTHEDKEVI<br>EEIAEYARKKGHFVVIVE |
| I-62 | MYQLIVVNAPKSLIREVVEEYGAKRERVYIPGPRVGAVIDVFYFDSKED<br>AEAVEERARAAGLEVFLIE |
| I-66 | AKKEIEELKKKAEEFYKKAEEEKKKFLENKDKFVKRIYYYAWRVGGV<br>WGITEDGEIISYGRANAYVAAAVLLEKKAEELKKKLEEEEK |
| I-73 | MEVILTNASLKKAKALAKELGVTYSVTYYYAWRVGGTYITLTGVTKEQ<br>AEKIQKELKVETYVVE |
| I-81 | MQVFIANVNAKTYKKCKEVAKKTGAKLTRTYYYAWRVGGIYITLEGV<br>DEETAKELEEFAKKEGNIVEIVK |

**Table S3. Cryo-EM data collection, data processing, and structure refinement statistics.**

| | NorA bound to miniprotein I-23 |
|---|---|
| **PDB ID** | **28VJ** |
| **EMDB ID** | **EMD-56885** |
| **Conformation** | **Outward-open** |
| **Data collection and processing** | |
| Magnification (x) | 105,000 |
| Voltage (kV) | 300 |
| Electron dose ($e^{-}/Å^2$) | 49.11 |
| Defocus range (μm) | 0.7 to 2.4 |
| Collection mode | Super-resolution |
| Pixel size (Å) | 0.4125 |
| Symmetry imposed | C1 |
| Initial number of particles | 6,691,933 |
| Final number of particles | 302,069 |
| Map sharpening *B* factor ($Å^2$) | 120.5 |
| Map resolution (Å)* | 2.79 |
| **Refinement** | |
| Non-hydrogen atoms | 3,167 |
| Protein residues | 412 |
| Mean B factor | |
| Protein ($Å^2$) | 69.06 |
| RMS deviations | |
| Bond lengths (Å) | 0.00 |
| Bond angles (°) | 0.45 |
| MolProbity score | 1.48 |
| Clash score | 5.27 |
| Rotamer outliers (%) | 1.80 |
| Ramachandran plot | |
| Favored (%) | 99.25 |
| Allowed (%) | 0.75 |
| Outliers (%) | 0.00 |
| Model resolution (Å)+ | 3.1 |

*Resolution determined by a FSC value of 0.143.

+Resolution determined between the model and the sharpened map by the FSC value of 0.5.
